# Phosphate transporter PHT1;1 as a key determinant of phosphorus acquisition in *Arabidopsis* natural accessions

**DOI:** 10.1101/2022.01.05.475091

**Authors:** Pei-Shan Chien, Ya-Ting Chao, Chia-Hui Chou, Yu-Ying Hsu, Su-Fen Chiang, Chih-Wei Tung, Tzyy-Jen Chiou

**Affiliations:** Agricultural Biotechnology Research Center, Academia Sinica, Taipei, Taiwan; Department of Agronomy, National Taiwan University, Taipei, Taiwan

## Abstract

To understand the genetic basis in governing phosphorus (P) acquisition, we performed genome-wide association studies (GWAS) on a diversity panel of *Arabidopsis thaliana* by two primary determinants of P acquisition, phosphate (Pi)-uptake activity and PHOSPHATE TRANSPORTER 1 (PHT1) protein abundance. Association mapping revealed a shared significant peak on chromosome 5 (Chr5) where the *PHT1;1*/*2*/*3* genes reside, suggesting a strong correlation between the regulation of Pi-uptake activity and PHT1 protein abundance. Genes encoding transcription factors, kinases, and a metalloprotease associated with both traits were also identified. Conditional GWAS followed by statistical analysis of genotype-dependent expression of *PHT1;1* and transcription activity assays revealed an epistatic interaction between *PHT1;1* and *MYB DOMAIN PROTEIN 52* (*MYB52*) on Chr1. Analyses of F1 hybrids generated by crossing two subgroups of natural accessions carrying specific SNPs associated with *PHT1;1* and *MYB52* further revealed the strong effects of potential variants on *PHT1;1* expression and Pi uptake activity. Notably, the soil P contents in *A. thaliana* habitats were found to coincide with *PHT1;1* haplotype, underscoring how fine-tuning of the activity of P acquisition by natural variants allows plants to adapt to their environments. This study sheds light on the complex regulation of P acquisition and offers a framework to systematically assess the effectiveness of GWAS approaches in the study of quantitative traits.

**One sentence summary:** Stepwise GWAS analyses reveal insights into the genetic basis in regulating phosphorus acquisition and associations between the phosphate transporter *PHT1;1* haplotype and *Arabidopsis* habitats.

## INTRODUCTION

Plants have evolved an array of strategies to sustain growth across diverse habitats where soil nutrients differ widely. Phosphate (Pi), the primary form of phosphorus (P) acquired by plants, is a limited resource because of its low solubility and mobility in soil (Theodorou and Plaxton, 1993; Raghothama, 1999). Modulation of P transport systems (Misson et al., 2004; Shin et al., 2004; Ai et al., 2009), root system architecture (Lynch, 2011; Pandey et al., 2013), secretion of root exudates (Wang et al., 2011), and reallocation and recycling of internal P (Raghothama, 1999) are known to be the major adaptive responses plants used to cope with low availability of P in the environments. Remarkable progress has been made in understanding the genetic and molecular mechanisms that control these responses in *Arabidopsis thaliana* (*A. thaliana*), particularly in the Columbia (Col-0) accession. For example, PHOSPHATE STARVATION RESPONSE 1 (PHR1) and PHR1-LIKE 1 (PHL1), which are MYB-coiled coil (CC) type transcriptional activators, are known to play major roles in the activation of P starvation-inducible genes (Rubio et al., 2001; Bustos et al., 2010; Sun et al., 2016). Members of the PHOSPHATE TRANSPORTER 1 (PHT1), PHT5, and PHOSPHATE 1 (PHO1) transporter families were found to mediate P acquisition, storage, and root-to-shoot translocation, respectively (Hamburger et al., 2002; Misson et al., 2004; Shin et al., 2004; Liu et al., 2016). The activity of PHT1 is increased by its trafficking to the plasma membrane through *PHOSPHATE TRANSPORTER TRAFFIC FACILITATOR1* (PHF1), but phosphorylation suppresses its trafficking (González et al., 2005; Bayle et al., 2011; Chen et al., 2015; Yang et al., 2020). Moreover, several regulatory modules of miRNAs and their target genes, such as miR399-PHO2 (a ubiquitin-conjugating E2 enzyme), miR827-Nitrogen Limitation Adaptation (NLA; a RING type ubiquitin E3 ligase), and miR156-SQUAMOSA PROMOTER BINDING PROTEIN-LIKE 3 (transcription factor) are intertwined to regulate Pi transport (Huang et al., 2013; Lin et al., 2013; Lei et al., 2016).

Many important agronomic traits are controlled by quantitative trait loci (QTLs). Identification of QTLs associated with a trait of interest and subsequent analyses of candidate genes often offer new strategies for crop improvement, including improvement of the use efficiency of nutrient ions. For example, using a QTL mapping strategy, (Reymond et al., 2006) identified *LOW PHOSPHATE ROOT1*, which explained 52% of the variation in primary root growth response to P deficiency in an *A. thaliana* Bay-0 × Shahdara recombinant inbred line population. In rice, *Phosphorus-Starvation Tolerance 1* (*PSTOL1*)/Phosphorus uptake1, which encodes a protein kinase, is an important gene/QTL that enhances grain yield by promoting root growth (Wissuwa et al., 2002; Gamuyao et al., 2012). The absence of *PSTOL1* from modern rice cultivars highlights the importance of exploring the diversity of natural variants for breeding crops with improved P acquisition ability (Gamuyao et al., 2012). Tremendous efforts have also been made to dissect the genetic basis of various P-related traits in other crops (Wang et al., 2010; Wang et al., 2010; Hufnagel et al., 2014; Gu et al., 2016). Despite the identification of several potential P use efficiency QTLs, only few have been functionally validated (Gamuyao et al., 2012; Zhang et al., 2014; Luo et al., 2019).

*A. thaliana,* which is grown as an inbred line and for which there is a wealth of natural accessions with publicly available whole genome sequences, is an attractive model for QTL identification using the genome-wide association (GWAS) approach (Alonso-Blanco and Koornneef, 2000; The 1001 Genomes Consortium, 2016). There have been several recent studies related to P deficiency in *A. thaliana* accessions, such as identification of candidate loci regulating responses to a combination of salinity and P starvation (Kawa et al., 2016), root growth under P and iron limitation (Bouain et al., 2019), and leaf anion homeostasis under different P supplies (El-Soda et al., 2019). These analyses highlight the power of using GWAS in *A. thaliana* to decipher new players in P stress responses. However, this approach may have some potential drawbacks. These include false positive associations resulting from cryptic population structure and kinships and false negatives caused by overcorrection of confounding factors or insufficient statistical power to detect rare alleles (Nordborg and Weigel, 2008). Thus, to validate the biological roles of detected loci, systematic follow-up analyses are necessary (Verslues et al., 2014; Bouain et al., 2019).

P acquisition is a key determinant of P use efficiency (PUE) in plants; however, little is known about how genotypic variations govern P acquisition in natural environments. In this study, we analyzed a set of ∼200 *A. thaliana* accessions to dissect the genetic architecture controlling Pi-uptake activity and the protein levels of PHT1 Pi transporters under moderately low Pi conditions (50 μM). We established a platform for analyses including phenotyping followed by conventional and conditional GWAS mapping and the association of phenotypes with selected F1 heterozygotes. We identified several genes involved in regulating P acquisition-related traits. Importantly, we found a connection between the *PHT1;1* haplotypes and P acquisition ability and the edaphic conditions in the growth sites of natural accessions. Our findings draw attention to genotypic variants of *A. thaliana* accessions that are adapted to the soil P content in their habitats and present information for potentially improving PUE in crops along with a strategic approach for GWAS to identify causal genes.

## RESULTS

### GWAS of Pi-uptake activity and PHT1;1*/*2*/*3 protein abundance under moderately low Pi

Two panels of *A. thaliana* accessions (see details in “MATERIALS AND METHODS”) representing worldwide diversity and used to develop a high-resolution haplotype map and whole-genome sequences (Cao et al., 2011; Horton et al., 2012) were investigated in this GWAS. To determine the conditions for phenotyping P-acquisition ability, we first examined the growth and Pi concentrations of Col-0 seedlings grown in the medium containing different concentrations of Pi (KH_2_PO_4_) ranging from 10 to 250 μM (Supplemental Figure S1). Because we intended to focus on the responses to Pi limitation while circumventing potential adverse effects caused by impaired growth, a moderately low concentration of Pi (50 μM) was chosen (Supplemental Figure S1A). The Pi concentrations of seedlings grown under this condition were significantly lower than those grown under Pi sufficient conditions (250 μM), but there was no significant difference in fresh weight (Supplemental Figure S1, B-E). Using this condition, we measured two important traits contributing to P-acquisition ability in 12-day-old seedlings. The first trait, root [^33^P]Pi-uptake activity, was measured by performing liquid scintillation counting of whole seedlings of 306 natural accessions. The second trait, protein abundance of PHT1 Pi transporters in the root, was quantified in 180 accessions by immunoblot analysis using an antibody that recognizes three Pi transporters (PHT1;1, PHT1;2, and PHT1;3) because of their high similarity.

After removing the accessions that showed inconsistent phenotypes across three to four biological replicates (15 accessions for uptake activity and two for protein abundance) and 27 accessions for uptake activity that could not be identified in the GWA-Portal database (Seren et al., 2012; The 1001 Genomes Consortium, 2016; Seren, 2018), 264 and 178 accessions were used for GWAS analysis of Pi-uptake activity and PHT1;1/2/3 protein level, respectively. These traits were normally distributed (Figure 1, A and B; Supplemental Table S1), suggesting that both are quantitative traits determined by multiple genes. Principal component analysis (PCA) of genotypic data revealed that PC1 only explained 3.15% and 2.9% of the total variation in the two populations analyzed for PHT1;1/2/3 protein level and Pi-uptake activity, respectively (Supplemental Figure S2), suggesting that the influence of population structure in detecting genotype-trait association is negligible. Therefore, we employed the accelerated mixed model (AMM) implemented in GWA-Portal (Seren et al., 2012) for our association analysis, in which kinship is considered to effectively control confounding signals due to relatedness, and the variance components are estimated to obtain the overall pseudo-heritability. Based on the phenotype and kinship matrix, we calculated the heritability and estimated how much variation was contributed by genotypic variation. Heritabilities of 0.388 and 0.735 were estimated for Pi-uptake activity and PHT1;1/2/3 protein level, respectively. As shown in the Manhattan plots, several major SNP clusters were associated with Pi-uptake activity, but only one cluster with −log_10_(*P*) > 6 was identified for PHT1;1/2/3 protein level (Figure 1, C and D). Compared with PHT1 protein abundance, it is conceivable that the degree of complexity in controlling Pi-uptake activity is much higher, which involves other factors in addition to Pi transporters, such as root architecture and root exudate. Remarkably, both traits shared a significant peak on chromosome 5 (Chr5), suggesting that these two traits are controlled by the same locus. Intriguingly, *PHT1;1*/*2*/*3* (*AT5G43350*/*AT5G43370*/*AT5G43360*) are located within the SNP peak on Chr5. This observation shows the effectiveness of our experimental design and the importance of PHT1 family members for natural variation in Pi-uptake activity.

**Figure 1.**
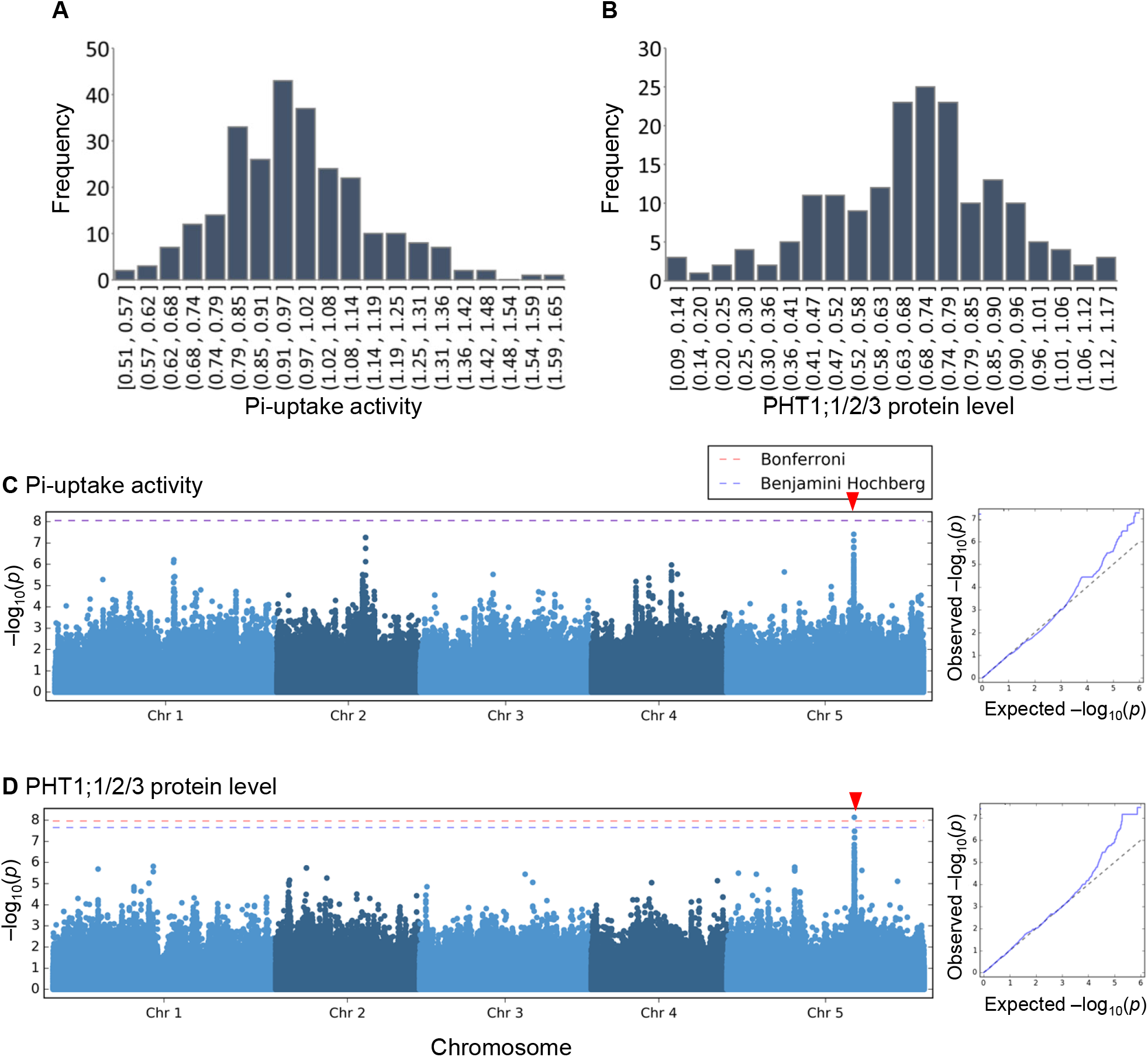
GWA analysis of two P-acquisition traits. Phenotypic distribution of raw values for Pi-uptake activity **(A)** and square root-transformed (SQRT) values for PHT1;1/2/3 protein abundance **(B)**. Manhattan plots of Pi-uptake activity **(C)** and PHT1;1/2/3 protein level **(D)**; the respective Q-Q plots are shown on the right. The analyses were conducted using GWA-Portal. Thresholds for Bonferroni (red dashed line) and Benjamini Hochberg (blue dashed line) correction provided by GWA-Portal are indicated. The two lines overlap in **(C)**.

### Identification of candidate genes associated with P-acquisition traits

To identify the candidate genes located in the regions adjacent to the significant peaks, we first defined significant SNPs and then searched for trait-associated candidate genes. The SNPs with a minor allele count (MAC) > 10 and a −log_10_(*P*) > 4, which was the cutoff based on the deviation of observed –log_10_(*P*) from expected – log_10_(*P*) in the quantile-quantile (Q-Q) plots (Figure 1, C and D, right panel), were selected. Using these criteria, 478 and 508 significant SNPs were identified for Pi-uptake activity and PHT1;1/2/3 protein abundance, respectively (Figure 2; Supplemental Table S2, A and B). As the decay of linkage disequilibrium (LD) in *A. thaliana* accessions is within 10 kb on average (Kim et al., 2007), we selected 462 and 282 protein-coding genes located within the 10-kb regions flanking the significant SNPs as potential candidates (Figure 2; Supplemental Table S3). Among them, 450 and 270 genes were unique for Pi-uptake activity and PHT1;1/2/3 protein level, respectively, and 12 genes were associated with both traits, including 4 Pi transporters, PHT1;1, PHT1;2, PHT1;3, and PHT1;6 (Figure 2; Supplemental Table S3).

**Figure 2.**
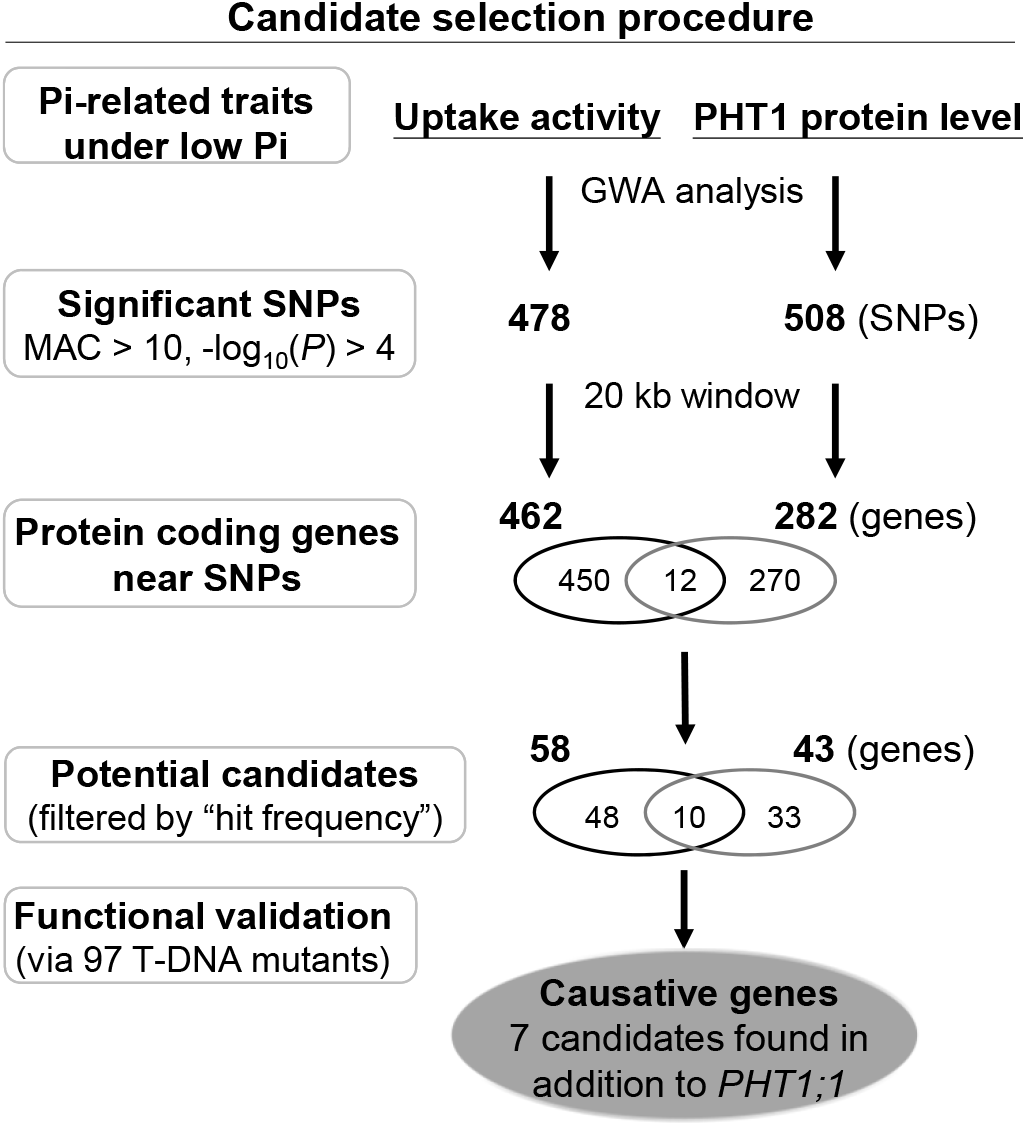
GWAS of two Pi-related traits and the step-by-step scoring system used for the selection of candidate genes. In addition to the well-known Pi transporters *PHT1;1* and *PHT1;2*, our approach identified seven candidate genes (e.g., transcription factors, a kinase, an oxidoreductase, and a protease) involved in P acquisition under low Pi.

To prioritize candidate genes for further functional assays, we assigned a value, denoted as “hit frequency”, based on the number of times a gene overlapped with the 10-kb region surrounding a significant SNP (Supplemental Figure S3). For instance, the hit frequency of *PHT1;1* was calculated to be 61 for uptake activity (Supplemental Table S3), meaning that any part of the *PHT1;1* gene body is within the 20-kb windows centered on 61 significant SNPs (Supplemental Figure S3). Genes were selected using the following criteria: (1) genes were correlated with either Pi-uptake activity or PHT1;1/2/3 protein level and had a hit frequency >10; or (2) genes were correlated with both traits and had a hit frequency >5. Accordingly, 81 genes associated with either Pi uptake activity (48 genes) or PHT1;1/2/3 protein level (33 genes) and 10 genes correlated with both were identified (Figure 2; Supplemental Table S3).

In line with the previous studies (Shin et al., 2004; Huang et al., 2013), *pht1*;*1* single mutant showed a significant and striking reduction in PHT1;1/2/3 protein level, Pi-uptake activity (Supplemental Figure S4), and Pi concentration (Figure 3) under a moderately low Pi condition (50 μM). In addition, we analyzed 97 available T-DNA insertional mutants covering 43 out of 91 candidate genes to evaluate their P-acquisition ability (Supplemental Table S4). We considered the genes promising candidates only when two independent allelic mutants showed consistent and significant changes. Using this criterion, we did not observe significant changes in the PHT1;1/2/3 protein level or short-term Pi-uptake activity for any of the candidate gene mutants (Supplemental Figure S4, A and B). However, when the Pi concentration of 12-day-old seedlings was analyzed, we found that mutations in seven of the genes, namely *AT1G17890* (*GDP-keto-6-deoxymannose 3,5-epimerase/4-reductase*; *GER2*), *AT1G17870* (*ETHYLENE-DEPENDENT GRAVITROPISM-DEFICIENT AND YELLOW-GREEN-LIKE 3*; *EGY3*), *AT1G17850* (Rhodanese/Cell cycle control phosphatase superfamily protein), *AT5G43330* (*CYTOSOLIC-NAD-DEPENDENT MALATE DEHYDROGENASE 2*; *C-NAD-MDH2*), *AT5G43320* (*CASEIN KINASE I-LIKE 8*; *CKL8*), *AT1G17950* (*MYB DOMAIN PROTEIN 52; MYB52*), and *AT2G27610* (tetratrico peptide repeat-like superfamily protein), showed a significant reduction (7%–23%) either in roots or shoots compared with the Col-0 wild-type (WT) control (Figure 3, A and B). Since Pi concentration represents a final consequence of various activities, including P acquisition, metabolism, and storage, we reasoned that these genes might participate in any of these activities despite a minor role in directly regulating Pi uptake. Nevertheless, it is also necessary to keep in mind that all these T-DNA insertional mutants are generated in a single accession, Col-0, and the complexity of naturally occurring variants from the *A. thaliana* genomic backgrounds in the diversity panel is not fully represented.

**Figure 3.**
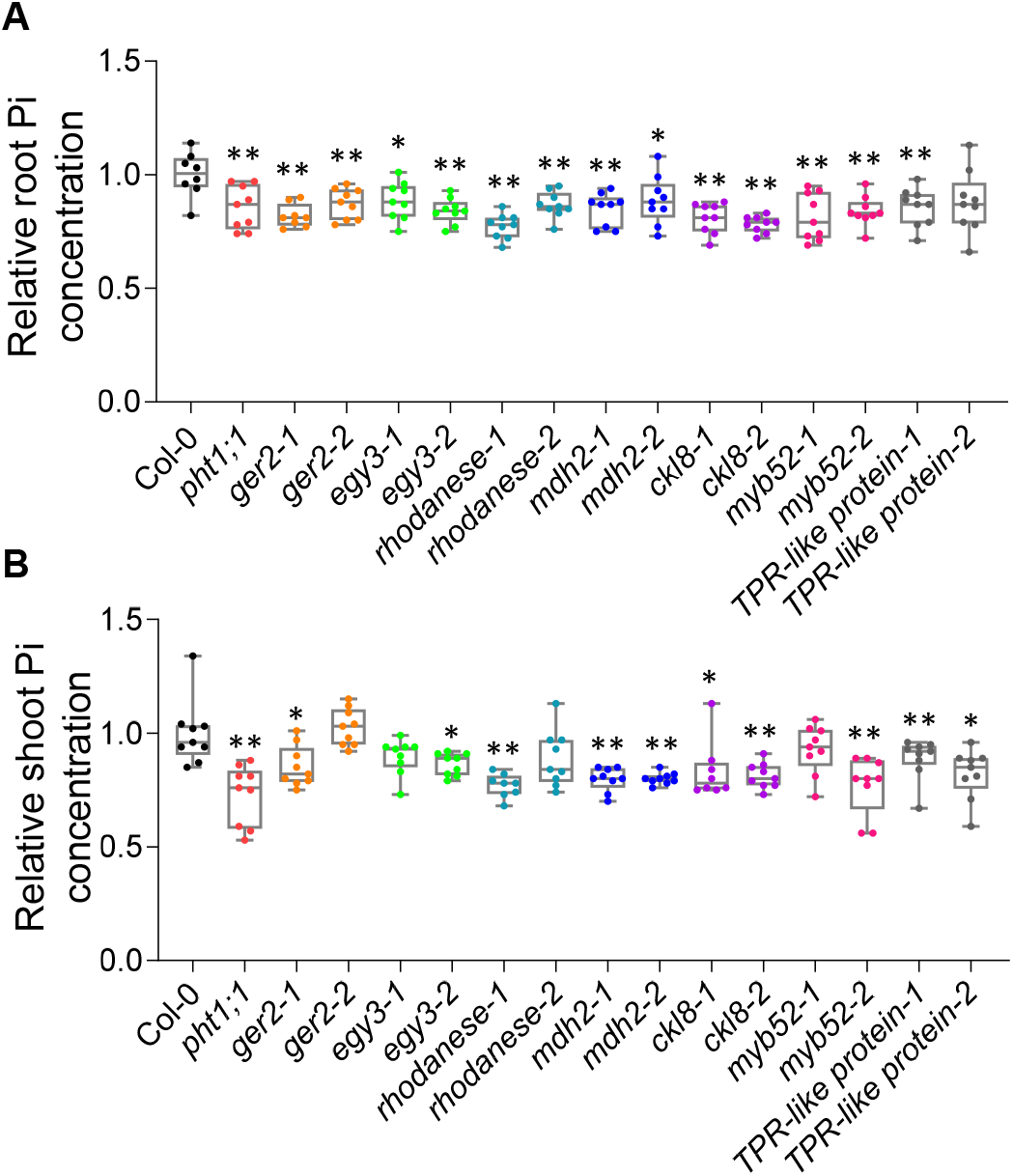
Analyses of Pi-related traits in the seedlings of T-DNA insertional mutants. Pi concentrations in mutants relative to Col-0 wild type in the roots **(A)** and shoots **(B)**. Twelve-day-old seedlings grown under moderately low Pi conditions (see “MATERIALS AND METHODS”) were sampled. The results are shown in a box and whisker diagram with individual data points superimposed. n = 9 biological repeats. The bars within the box plots represent the 25th and 75th percentiles, the center line represents the median, and the whiskers show the minimum and maximum values. Significant differences between mutants and Col-0 were determined by two-sided Student’s *t*-test, and *P*-values were adjusted for multiple testing using Benjamini– Hochberg correction (* Adjusted *P* < 0.05; ** Adjusted *P* < 0.01). Insertion mutants for the following genes were tested: *GER2* (*AT1G17890*); *EGY3*(*AT1G17870*); Rhodanese (*AT1G17850*); *C-NAD-MDH2* (*AT5G43330*); *CKL8*(*AT5G43320*); *MYB52* (*AT1G17950*), and tetratrico peptide repeat (TPR)-like superfamily protein (*AT2G27610*).

We then examined the RNA levels of selected genes in the roots and shoots of Col-0 under sufficient Pi (1 mM KH_2_PO_4_) and low Pi (50 μM KH_2_PO_4_) conditions (Supplemental Figure S5). Under sufficient Pi conditions, the RNA levels of most candidate genes were higher in the shoots than in the roots except that the *MYB52* transcription factor was mainly expressed in the roots and *GER2*, which encodes an enzyme involved in the biosynthesis of GDP-fucose, was expressed equally in both tissues. In response to low Pi conditions, only the expression level of *GER2* in the shoots was reduced slightly (0.8 fold). None of the other candidate genes were responsive to low Pi in the roots.

### Epistatic interaction between PHT1;1 and MYB52

Because there was only one prominent SNP peak on Chr5 associated with PHT1;1/2/3 protein level (Figure 1D), to identify additional loci, we performed conditional GWAS implemented in GWAPP (GWAS-Web-App, http://gwas.gmi.oeaw.ac.at; Seren et al., 2012), a predecessor of GWA-Portal. With that, the effect from a specific causal SNP could be included in the model to increase the power to detect other causal markers in a structured dataset when using AMM (Seren et al., 2012). This approach has been successfully applied in flowering phenotype and other GWAS studies (Atwell et al., 2010; Yang et al., 2012). After masking the effect of the most significant SNP on Chr5 (17,400,077 bp; the allele is C or T) (Figure 4A), a novel SNP peak on Chr1 became evident (Figure 4B). This result implies a biological interaction between the genes where the significant SNPs on Chr1 and Chr5 are located. The most significant SNP is located on *HOMEODOMAIN GLABROUS 12* (*HDG12*; *AT1G17920*), which belonged to an HD-ZIP IV family. However, the T-DNA mutants of *HDG12* did not show differences in Pi concentrations from WT controls. We thus designated the second significant SNP on Chr1 located at 186 bp downstream of the 3’-UTR of *MYB52* (*AT1g17950*), whose association with Pi accumulation in roots was identified in our analysis (Figure 3A), as SNP locus 1 (6,179,475 bp; the allele is A or T; Supplemental Figure S6) and the one on Chr5 located in the third exon of *PHT1;1* as SNP locus 2 (Supplemental Figure S7A). There are four possible allele combinations, {A,C}, {A,T}, {T,C} and {T,T}, of these two significant SNP loci.

**Figure 4.**
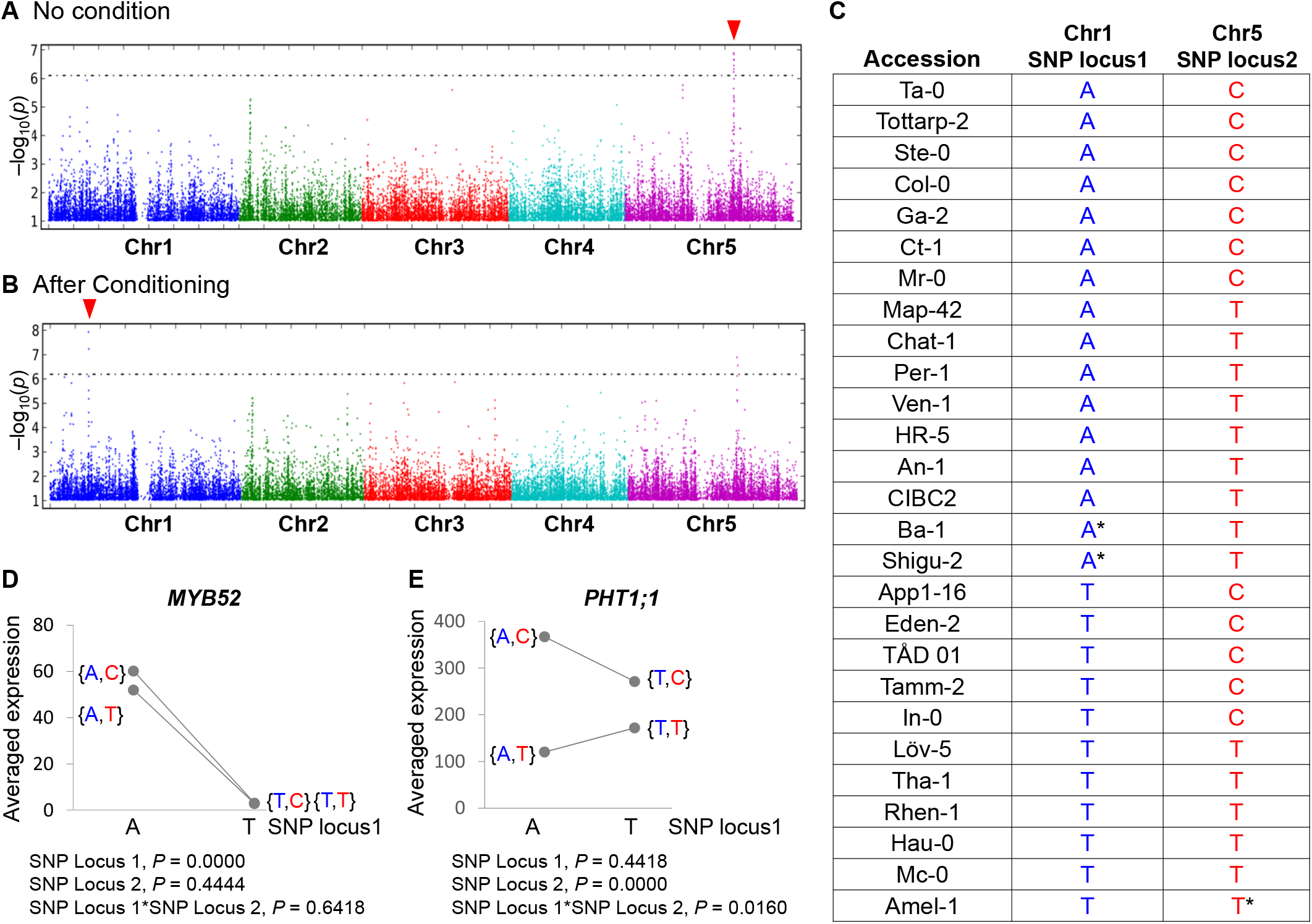
Conditional GWAS and statistical analyses reveal an association between *MYB52* and *PHT1;1.* Manhattan plots of GWAS analyses performed using AMM **(A**, no conditioning) or modified AMM in which the most significant SNP located at Chr5 (the arrowhead in **(A)**) was included as a cofactor **(B**, conditioning) in GWAPP. The 5% false discovery rate threshold is plotted as a dashed horizontal line. **(C)** List of candidate accessions used for examination of the epistatic relationship between Chr1 and Chr5. Asterisks (*) indicate the SNPs that were corrected after re-sequencing. Analyses of the relationship between SNP locus 1 and SNP locus 2 by two-way ANOVA based on average gene expression levels of *MYB52* (2^-△CT^ x 1000) **(D)** and *PHT1;1* (2^-△CT^ x 100) **(E)**. The letters in blue and red shown in **(C)**, **(D)**, and **(E)** indicate SNP locus 1 and SNP locus 2, respectively.

Next, we selected accessions based on the four allele combinations to test the potential interaction between these two SNP loci. Because the population in four different genotype combinations among the 178 accessions analyzed is varied, based on the availability of genome sequences in *Arabidopsis* 1001 Genome (https://tools.1001genomes.org/vcfsubset/#select_strains), we randomly selected 5-9 accessions from each allele combination as representatives (6 out of 8 {T,T}-accessions, 9 out of 63 {A,T}-accessions, 7 out of 92 {A,C}-accessions and 5 out of 15 {T,C}-accessions) (Figure 4C). We then evaluated the expressions of 6 annotated candidate genes surrounding the SNP locus 1 (Supplemental Figure S6A; the lower panel) and *PHT1;1*/*2*/*3* genes (SNP locus 2) in each accession. Among the genes surrounding SNP locus 1, only *MYB52* showed a clear difference in expression between the two allelic variants at SNP locus 1 (Supplemental Figure S8B). As revealed by two-way ANOVA, *MYB52* expression was significantly higher in the accessions with allele “A” than in those with allele “T” at SNP locus 1 (*P* < 1.00E-04), regardless of the allele at SNP locus 2 (Figure 4D). Intriguingly, a significant allelic interaction between SNP locus 1 and SNP locus 2 was observed for the expression of *PHT1;1*(*P* = 1.60E-02) (Figure 4E). These observations highlight the existence of an interaction between *MYB52* and *PHT1;1*. On the other hand, there was no significant allelic interaction between SNP locus 1 and 2 for *PHT1;2* (*P* = 4.16E-01) or *PHT1;3* (*P* = 6.99E-01) (Supplemental Figure S8, C and D).

Because MYB52 was annotated as an R2R3-MYB transcription factor (Dubos et al., 2010), we hypothesized that there is an epistatic interaction between *MYB52* and *PHT1;1* resulting from MYB52-dependent transcriptional regulation of *PHT1;1*. To examine this possibility, we co-expressed Col-0-MYB52 as an effector with a luciferase reporter driven by the promoter of Col-0-*PHT1;1* in protoplasts isolated from the leaves of Col-0 (Figure 5A). In supporting our hypothesis, MYB52 positively regulated *PHT1;1* promoter activity (1.8 fold of the vector control) (Figure 5B). However, the activation was not as high as the major Pi regulator PHR1 (5.4 fold of the vector control) (Figure 5B). On the other hand, we were unable to detect a physical interaction between MYB52 and its potential binding sites (MBSII) in the promoter of *PHT1;1* by yeast one-hybrid analysis despite the occurrence of positive interaction between PHR1 and the PHR1-binding sequences (P1BS) in the *PHT1;1* promoter (Supplemental Figure S9). Because MYB is one of the transcription factor families that can act as homo-or heterodimers to regulate distinct cellular processes (Pireyre and Burow, 2015), MYB52-mediated transcriptional activation of *PHT1;1* may require other factors.

**Figure 5.**
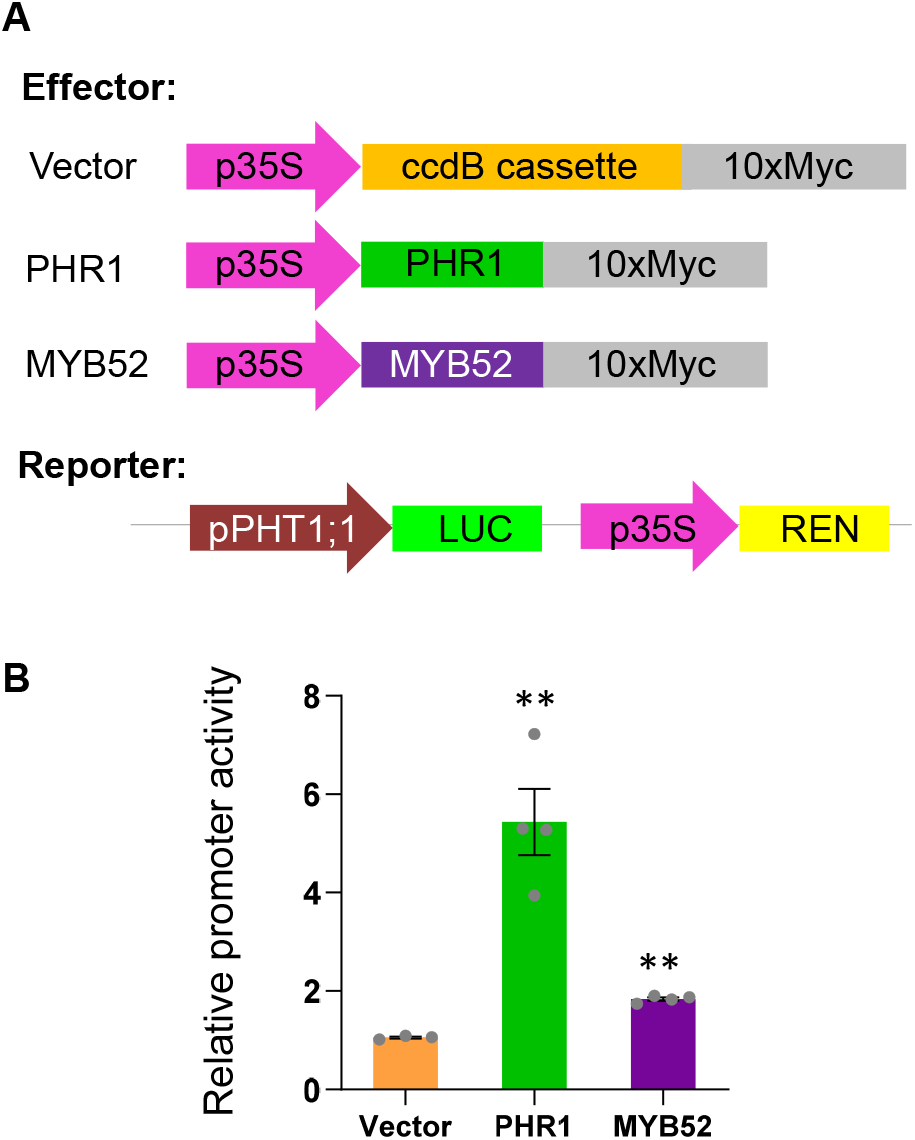
MYB52 regulates the promoter activity of *PHT1;1.* **(A)** A schematic illustration of constructs used for the analysis of *PHT1;1* promoter activity in a dual-luciferase assay. **(B)** *PHT1;1* promoter activity was defined as the ratio of LUC:REN (firefly:renilla). Transformation with the empty vector served as a negative control. Data are presented as mean ± SE (n = 4 biologically independent samples). The significance of differences was determined by two-sided Student’s *t*-test (** *P* < 0.01).

### {A,C} genotype is correlated to the higher PHT1;1 expression and the superior Pi uptake activity

Given that the accessions with the {A,C} allele combination exhibited the highest *PHT1;1* gene expression among the four combinations (Figure 4E), we were curious how P acquisition is affected when one genomic copy of the {A,T} allele is replaced by one copy of the allele from the {A,C} genetic background. Thus, we crossed {A,T} accessions (Cnt-1, Shigu-2, An-1, Alst-1, CIBC2, Ba-1, and Ws) or {A,C} accessions (Paw-3, Ciste-1, Voeran-1, Sei-0, Ta-0, and Ga-2) to Col-0, which is also {A,C} (Table 1). Although we have yet to dissect potential factors at other genomic loci that may also contribute to the phenotype, we observed significantly increased *PHT1;1* expression and Pi-uptake activity relative to the maternal {A,T} accessions when one copy of the {A,C} genome was introduced into the {A,T} background. For instance, F1 plants from Cnt-1 X Col-0, which are {A,T}X{A,C} hybrids, showed a 7.19-fold increase in *PHT1;1* gene expression and a 1.57-fold increase in Pi uptake activity when compared with the maternal line Cnt-1. This phenomenon was not observed in the {A,C}X{A,C} F1 hybrid (Table 1). We surmise that the varied P acquisition ability in the {A,C}X{A,C} hybrids may cause by heterosis or hybrid weakness. These results showed that introduction of one copy of the {A,C} genome can improve the P acquisition of the {A,T} accessions, probably through the introduction of functional *PHT1;1*.

**Table 1.** P-acquisition phenotypes of F1 hybrids.

| Accession ID | Genotype | *PHT1;1* expression<sup>a</sup> |  |  | Uptake activity<sup>b</sup> |  |  |
| --- | --- | --- | --- | --- | --- | --- | --- |
|  |  | Maternal line | F1<sup>d</sup><br>(X Col-0) | F1<sup>d</sup><br>(X *pht1;1*) | Maternal line | F1<sup>d</sup><br>(X Col-0) | F1<sup>d</sup><br>(X *pht1;1*) |
| An-1 | {A,T} | 12.3 ± 3.7 | 18.4 ± 3.8 | 5.4 ± 0.3 | 1.4 ± 0.1 | 1.7 ± 0.1* | 1.4 ± 0.1 |
| Ba-1 | {A,T} | 29.0 ± 6.3 | 52.4 ± 1.9* | 11.2 ± 1.0 | 1.6 ± 0.0 | 1.9 ± 0.1* | 1.3 ± 0.1* |
| CIBC2 | {A,T} | 8.1 ± 1.2 | 22.8 ± 1.6** | 8.8 ± 1.7 | 1.6 ± 0.0 | 1.8 ± 0.1* | 1.7 ± 0.2 |
| Cnt-1 | {A,T} | 4.8 ± 1.6 | 34.5 ± 5.3** | 4.6 ± 1.1 | 1.0 ± 0.0 | 1.6 ± 0.0** | 1.3 ± 0.1** |
| Shigu-2 | {A,T} | 21.5 ± 2.2 | 38.5 ± 3.3** | 12.4 ± 1.9* | 1.0 ± 0.1 | 1.7 ± 0.0** | 1.4 ± 0.1* |
| Ws | {A,T} | 20.0 ± 1.8 | 25.3 ± 5.4 | 15.6 ± 2.9 | 1.7 ± 0.1 | 1.7 ± 0.1 | 1.4 ± 0.1* |
| **Col-0** | {A,C} | 43.9 ± 6.0 | — | 23.5 ± 4.3* | 1.4 ± 0.1 | — | 1.2 ± 0.1* |
| Alst-1 | {A,C} | 9.3 ± 1.1 | 47.6 ± 6.5** | 16.1 ± 2.9 | 1.5 ± 0.1 | 2.0 ± 0.1** | 1.9 ± 0.1* |
| Ciste-1 | {A,C} | 68.5 ± 9.8 | 72.8 ± 9.6 | 29.1 ± 10.6* | 1.9 ± 0.1 | 1.7 ± 0.1 | 1.6 ± 0.1 |
| Ga-2 | {A,C} | 104.4 ± 22.8 | 87.1 ± 8.9 | 41.9 ± 4.7* | 2.3 ± 0.1 | 2.1 ± 0.2 | 1.4 ± 0.1** |
| Paw-3 | {A,C} | 62.6 ± 5.0 | 47.3 ± 1.7* | 26.7 ± 7.3* | 1.5 ± 0.1 | 1.8 ± 0.1 | 1.4 ± 0.1 |
| Sei-0 | {A,C} | 83.9 ± 6.7 | 47.9 ± 6.6* | 33.4 ± 3.3** | 2.1 ± 0.1 | 2.0 ± 0.1 | 1.7 ± 0.1* |
| Ta-0 | {A,C} | 31.8 ± 3.2 | 50.4 ± 6.0* | 23.7 ± 2.9 | 2.3 ± 0.1 | 1.9 ± 0.1 | 1.6 ± 0.1** |
| Voeran-1 | {A,C} | 58.8 ± 6.5 | 31.8 ± 3.5* | 25.6 ± 4.5* | 1.9 ± 0.0 | 1.9 ± 0.1 | 1.7 ± 0.1 |
| ***pht1;1*** | {A,C} | 0.0 ± 0.0 | — | — | 1.0 ± 0.1 | — | — |

The indicated trait value was averaged from three to four biological repeats and the value is shown as average ± SE.
Asterisks mark values significantly different between the maternal line and F1 hybrids (\* $P < 0.05$ ; \*\* $P$ $< 0.01$ , Student's $t$ -test).
<sup>a</sup> indicates the relative *PHT1;1* gene expression value ( $2^{-\Delta CT} \times 10$ ).
<sup>b</sup> indicates the relative Pi-uptake activity from each accession normalized to the level from the *pht1;1* mutant.
<sup>c</sup> Ratio indicates the average Pi trait value of F1 hybrids divided by that of inbred lines.
<sup>d</sup> F1 indicates maternal line x Col-0.
—, not applicable.

We also site-by-site generated another F1 population by crossing a homozygous *pht1*;*1* knockout mutant (Col-0 background) with various {A,T}-and {A,C}-accessions, the accessions selected above (Table 1). We found that the *PHT1*;*1* expression was rescued partially in both {A,T}X*pht1*;*1* and the {A,C}X*pht1*;*1* hybrids (Table 1), indicating that *PHT1*;*1* introduced from either {A,T}-or {A,C}-genotype is functional. The partial complementation may result from one copy of functional *PHT1*;*1* (i.e. *pht1*;*1*^Col-0^/*PHT1*;*1*^{A,T}^ or *pht1*;*1*^Col-0^/*PHT1*;*1*^{A,C}^). It is interesting to note that the *PHT1*;*1* expression in the {A,C}X*pht1*;*1* hybrid was higher than that in the {A,T}X*pht1*;*1* hybrid. However, the difference in the Pi-uptake activity between these two hybrids was not evident, implying other effects from unidentified loci complex with *PHT1*;*1* expression.

The results of F1 analysis support the genetic interaction between SNP locus 1 and locus 2: (1) *PHT1*;*1* gene expression increased significantly in the {A,C}X*pht1*;*1* hybrids than in the {A,T}X*pht1*;*1* hybrids, and (2) the performance for P acquisition is significantly increased in the {A,T}X{A,C} hybrids but not the {A,C}X{A,C} hybrids. Conclusively, {A,C} genotype is correlated to the higher *PHT1*;*1* expression and the superior Pi uptake activity.

Because PHT1;1 is a major determinant of Pi-uptake activity and its expression is highly dependent on the allelic variation at SNP locus 2, i.e., the accessions with allele “C” had significantly higher transcript levels than those with allele “T” (Figure 4E), we divided the *A. thaliana* diversity panel into two populations based on the allelic variation at SNP locus 2 (either “C” or “T”) and searched for additional loci controlling Pi-uptake activity independent of PHT1;1. Using these two artificial populations composed of 153 accessions with allele “C” and 111 accessions with allele “T”, we conducted GWAS for Pi-uptake activity. Although a few SNP peaks were observed in the Manhattan plots, none were significant (-log_10_(*P*) > 6, based on the threshold used in Figure 1, C and D) (Supplemental Figure S10). Therefore, Pi-uptake activity is mainly controlled by SNP locus 2 on Chr5. The effect of other QTLs may not be substantial enough to be detected by this approach.

### The PHT1 haplotype as a hallmark linking P-acquisition traits and the soil P contents in A. thaliana habitats

Because SNP locus 2 is a synonymous SNP, we searched for non-synonymous SNPs and found four nearly consecutive significant SNPs in the second exon of *PHT1;1* (Chr5: 17,401,478, 17,401,472, 17,401,471, and 17,401,466 bp) denoted as SNP3, SNP4, SNP5, and SNP6, respectively (Supplemental Figure S7B). These SNPs are in high LD (r^2^ = 0.99 in the group of accessions assayed for PHT1;1/2/3 protein level, Supplemental Table S5A) and explain 9.11% and 10.25% of the genotypic variance in Pi uptake activity and PHT1;1/2/3 protein level, respectively (Supplemental Table S2, A and B). These SNPs code for two forms of the PHT1;1 protein, the Col-0 type Asn-Pro-Glu-Ser-Ala (NPESA) and the non-Col-0 type Asp-Pro-Thr-Ser-Pro (DPTSP) (Supplemental Figure S7B). To investigate whether these non-synonymous SNPs could alter P-acquisition activity, we first inspected the subcellular localization of the proteins encoded by these SNP alleles and found no difference in their plasma membrane localization (Supplemental Figure S11A). Then, we examined the protein level and Pi-uptake activity in *pht1;1* mutants expressing PHT1;1^NPESA^ or PHT1;1^DPTSP^ driven by the native *PHT1;1* promoter (Supplementary Figure S11, B and C). Despite expression variation among independent transgenic lines, in general, both transgenes PHT1;1^NPESA^ and PHT1;1^DPTSP^ could restore the phenotypes (PHT1;1/2/3 protein level and Pi-uptake activity) of the *pht1;1* mutant to the WT level (Supplemental Figure S11, B and C). However, no consistent significant differences were observed between plants expressing these transgenes, suggesting that the corresponding nucleotides may serve as markers rather than causative SNPs.

SNP locus 2 is in high LD with SNP3/4/5/6 (r^2^ = 0.67 pairwise in the group of accessions assayed for PHT1 protein level, Supplemental Table S5A). The combination of these five SNPs results in four haplotypes, GACCT, GACCC, AGAGT, and AGAGC (Supplemental Figure S7C), in which GACCT (105 and 68 accessions assayed for Pi-uptake activity and PHT1;1/2/3 protein level, respectively) and AGAGC (139 and 94 accessions assayed for Pi-uptake activity and PHT1;1/2/3 protein level, respectively) are the two major haplotypes. It is of interest to note that the accessions with the “AGAGC” haplotype showed significantly higher Pi-uptake activities (*P* = 3.39E-08) and PHT1;1/2/3 protein levels (*P* = 3.64E-08) than those with the “GACCT” haplotype (Figure 6A and 6B). Notably, when we located the habitats of these natural accessions in Europe, we found a significant association (*P* = 1.47E-02) between these two haplotypes and the edaphic topsoil conditions (Tóth et al., 2014) (Figure 6C). The high Pi-uptake accessions (with the “AGAGC” haplotype) tended to be collected from growing sites containing low soil P while the low Pi-uptake accessions (with the “GACCT” haplotype) were collected from high soil P sites (Figure 6D). However, the relatively weak correlation with Pi levels suggests that soil P is not the only factor to impose selection pressure on *A. thaliana* populations. In summary, our analyses revealed the importance of the *PHT1;1* haplotype in connecting the P-acquisition activities of natural *A. thaliana* accessions to the soil P contents in their habitats.

**Figure 6.**
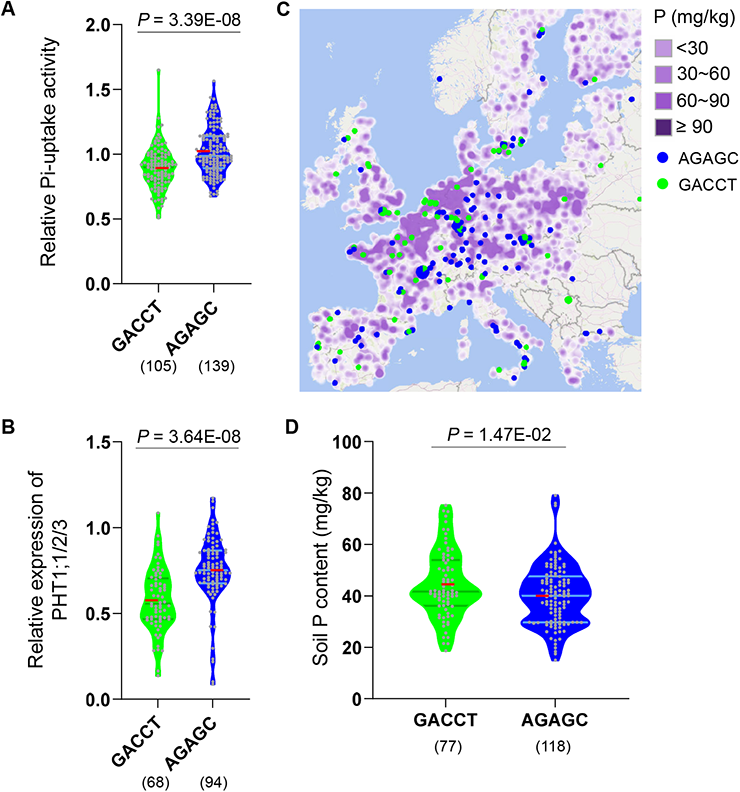
*PHT1;1* haplotype connects the P-acquisition phenotypes of *A. thaliana* accessions to the soil P contents in their habitats. Comparison of Pi-uptake activity **(A)** and PHT1;1/2/3 protein level **(B)** between accessions with different *PHT1;1* haplotypes. Each gray dot represents an individual accession. **(C)** Distribution of two different genotypic accessions and soil P-contents (in purple) on a map of the European Union. Symbols in green are accessions carrying the “GACCT” haplotype, while symbols in blue are those with “AGAGC”. **(D)** Comparison of soil P content for two types of accessions: those with the “GACCT” haplotype and those with the “AGAGC” haplotype. The gray lines within violin plots represent the 25th percentile, median, and 75th percentile. The red lines indicate the means of the quantified phenotypes. The *P*-value determined by two-sided Student’s *t*-test is indicated on the top of each plot. Values in parentheses shown below the SNPs indicate the numbers of accessions assayed.

## DISCUSSION

### PHT1;1 is a key determinant of P-acquisition activity as evidenced by analysis of A. thaliana natural accessions

In this study, we employed GWAS to analyze the genetic architecture of two major traits representing the capability to acquire P. Figure 7 illustrates the critical steps of our systematic approach. We identified a significant SNP peak for both traits on Chr5 where the Pi transporters PHT1;1, PHT1;2, and PHT1;3 are located (Figure 1), highlighting the importance of these Pi transporter genes in regulating their own protein abundance as well as the Pi-uptake activity of whole plants. In agreement with a very recent report (Yi et al., 2021), the *PHT1* was also identified as one of the crucial loci conferring natural variations in low Pi tolerance using five Pi starvation response traits. These findings reinforce the importance of the PHT1 family for P acquisition from the perspective of genetic variations in a natural population. Nevertheless, (Sakuraba et al., 2018) employed ^33^Pi radioactive imaging to analyze 200 natural accessions of *A. thaliana* but did not identify any QTL markers for P-acquisition ability. This is probably because the sensitivity of the imaging system was lower than that of the scintillation counter employed in this study.

**Figure 7.**
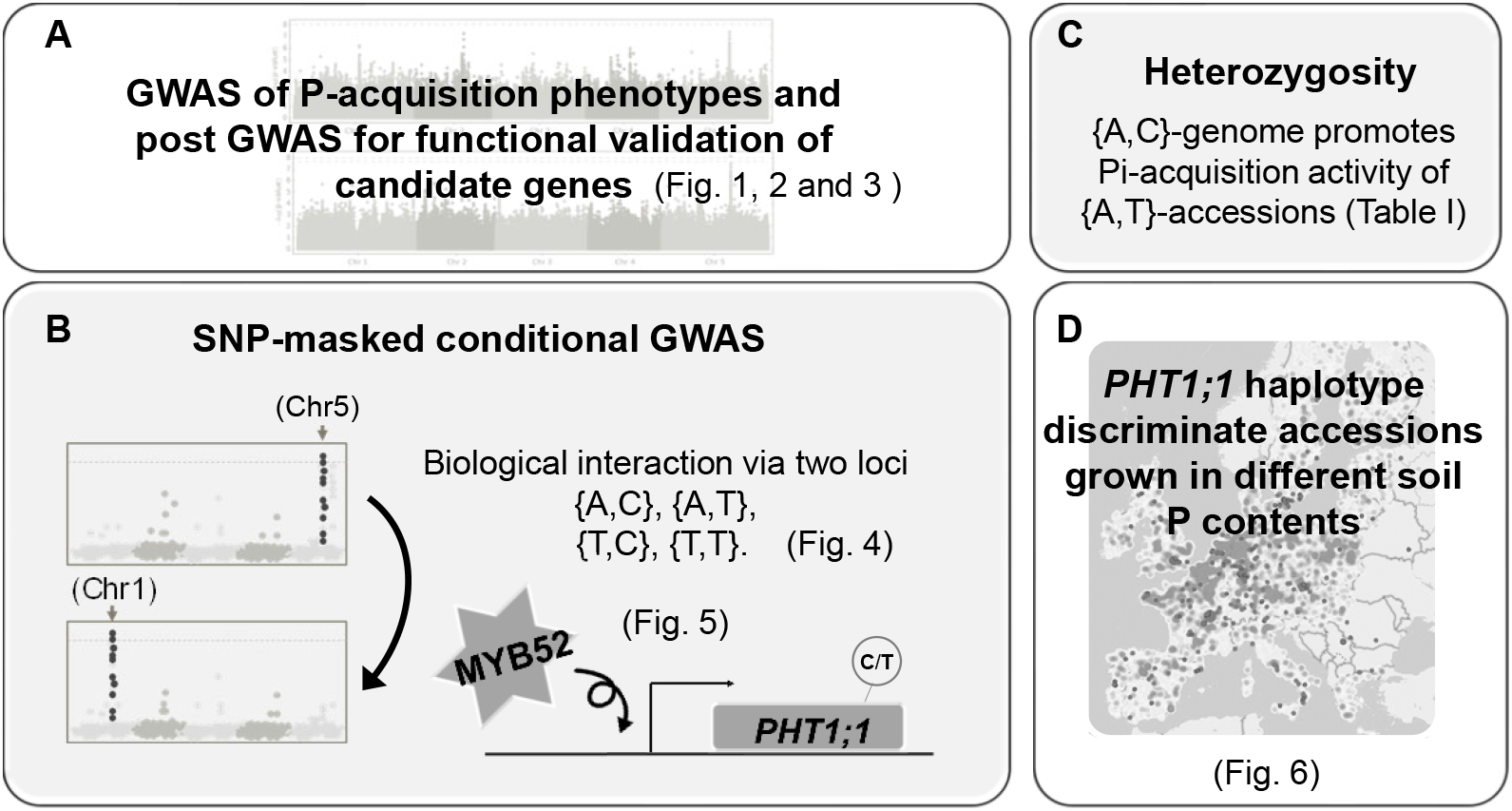
Illustration of our GWAS approach. **(A)** GWAS of two Pi-related traits. Our approach identified the well-known PHT1 Pi transporters and an additional seven genes potentially involved in regulating P acquisition under low Pi. **(B)** Conditional GWAS of PHT1;1/2/3 protein level revealed a biological interaction between SNP locus 1 and locus 2 and an MYB52-PHT1;1 regulatory module. **(C)** The results of heterozygosity analysis revealed improvement of Pi phenotypes by introduction of the {A,C} genome into {A,T} lines. **(D)** Different *PHT1;1* haplotypes discriminate soil P contents in the *A. thaliana* habitats.

Among PHT1;1, PHT1;2 and PHT1;3, PHT1;1 plays the dominant role, possibly because of its relatively high expression level (Supplemental Figure S8, C and D). Indeed, unlike the *pht1;1* mutant, the *pht1;2* and *pht1;3* mutants showed no reduction in the Pi concentration compared with the WT Col-0 (Huang et al., 2013). Yi et al. (2021) hypothesized this might result from gene duplication followed by a strong purifying selection to adapt to the environment. Our analyses clarified the difference in regulating the transcription level of these *PHT1* homologs among *A. thaliana* natural accessions, revealing that *PHT1;1* is controlled by MYB52, but not *PHT1;2* or *PHT1;3* (Figure 4; Supplemental Figure S8). Importantly, we provide evidence to connect the *PHT1*;*1* haplotype of natural accessions to the soil P contents in their habitats.

### MYB52 regulates PHT1;1 expression and Pi accumulation

Expression of *A. thaliana PHT1;1* was previously reported to be positively regulated by PHR1 (Rubio et al., 2001), WRKY DNA-BINDING PROTEIN 75 (WRKY75) (Devaiah et al., 2007), WRKY45 (Wang et al., 2014), WRKY42 (Su et al., 2015), phytochrome interacting factors PIF4/PIF5 and ELONGATED HYPOCOTYL 5 (Sakuraba et al., 2018), and NITRATE-INDUCIBLE, GARP-TYPE TRANSCRIPTIONAL REPRESSOR 1/HYPERSENSITIVE TO LOW PI-ELICITED PRIMARY ROOT SHORTENING 1.2 (Wang et al., 2020). Our study employed conditional GWAS followed by epistasis analysis and transcription activity assays and identified MYB52 as an additional positive regulator of *PHT1;1* expression (Figures 4 and 5). The observation of reduced shoot Pi concentration in *myb52* mutants (17.5% reduction compared with WT, Figure 3A) further supports this notion. However, we did not observe significant changes in PHT1;1/2/3 protein level or short-term Pi-uptake activity in the *myb52* mutants (Supplemental Figure S4). We think the reason for this is that the T-DNA insertional mutants generated in a single Col-0 background may not fully represent the complexity of genetic variations found in the diversity panel of *A. thaliana* natural accessions analyzed in this work. In addition, in the *myb52* single mutant, other positive transcriptional activators may compensate *PHT1*;*1* expression for the loss of *MYB52*.

So far, only a limited number of studies have explored the function of MYB52, and these studies have focused on its potential roles in regulating ABA-mediated drought tolerance and cell wall thickening (Zhong et al., 2008; Park et al., 2011). Although cell wall modification is associated with the change of root architecture and P re-utilization (Lin et al., 2011; Zhu et al., 2015; Naumann et al., 2019), involvement of MYB52 in these processes requires further investigation. Several other R2R3 MYB transcription factors were identified to be upregulated by P deficiency and involved in P acquisition or P-starvation responses. For example, *A. thaliana* MYB2 and MYB62 regulate P-starvation responses via an miR399-mediated pathway and gibberellic acid biosynthesis, respectively (Devaiah et al., 2009; Baek et al., 2013). In rice, overexpression of *OsMYB5P* and *OsMYB2P-1* could improve P acquisition by enhancing the expression of Pi transporters (Dai et al., 2012; Yang et al., 2018). Unlike these MYB transcription factors, the expression of *MYB52* does not respond to changes of external Pi supply (Supplemental Figure S5), underlining the value of GWAS, in which analysis is independent of gene expression.

### PHT1;1 haplotype connects the P-acquisition phenotypes of A. thaliana accessions to the soil P contents in their habitats

Based on the epistatic analysis of SNP locus 1 and SNP locus 2, the strength of the regulation of *PHT1;1* gene expression could be differentiated by allele combination, with the highest *PHT1;1* expression occurring in accessions carrying {A,C} and the lowest in those having {A,T} (Figure 4E). Moreover, analysis of heterozygotes (Table 1) revealed the potential for increasing P acquisition ability by introduction of the {A,C} allele. These observations highlighted the importance of SNPs within (or near) the *PHT1;1*gene body in regulating PHT1 protein abundance and P acquisition. Notably, we found that two groups of accessions, discriminated by *PHT1;1* haplotype (SNP3/4/5/6/locus 2), showed different P-acquisition phenotypes and different soil P contents in their habitats. As shown in Figure 6, the“AGAGC” haplotype is found in high-activity accessions preferentially collected from low-P soil. In contrast, the “GACCT” haplotype corresponds to low-activity accessions preferentially collected from high-P soil. This result suggests that soil P level imposes selection pressure on *A. thaliana* populations. The accessions with the “AGAGC” haplotype may adapt better to poor P soil than those with the “GACCT” haplotype because of improved ability to acquire P. To our knowledge, this is the first report to show the association between P-related traits and soil P content.

### Discovery of additional regulators of P acquisition

Apart from *PHT1;1*/*2*/*3*, we did not identify other well-known genes that regulate P-acquisition activity, such as *PHR1*, *PHO2*, *NLA*, and *PHF1* (Rubio et al., 2001; González et al., 2005; Huang et al., 2013; Lin et al., 2013). It is not surprising that GWAS fails to identify known genes. In a similar attempt, *PHT1* loci is the only well-known Pi regulatory cluster identified in the GWAS analysis, even though five P-acquisition-related traits were analyzed (Yi et al., 2021). The detection power is affected by many factors, such as allele frequency of variants, population size, and demographic history of the analyzed trait (Korte and Farlow, 2013). Nonetheless, we identified several new loci potentially involved in P acquisition (Figure 3). For example, C-NAD-MDH2 can convert oxaloacetate to malate, secreting into the rhizosphere to liberate Pi from the soil (Ryan et al., 2001). CKL8, a Ser/Thr kinase, was reported to modify the activity of ACC synthase 5 (Tan and Xue, 2014) and may regulate root elongation via ethylene under Pi-deprivation (Ma et al., 2003). EGY3 was classified as a site-2 protease-like putative metalloprotease implicated in endoplasmic reticulum stress (Iwata et al., 2009; Iwata et al., 2017), which could trigger Pi-dependent activation of autophagy (Naumann et al., 2019). The roles of CKL8 and EGY3 in P acquisition require further investigation. It is of interest to note that the expression of these candidate genes is not responsive to changes of Pi supply (Supplemental Figure S5), making it impossible to identify them by transcriptomic analysis (Misson et al., 2005; Liu et al., 2016). Thus, GWAS analysis complements transcriptomic analysis.

It is always challenging to validate the result of GWAS. Even with sufficient statistical power to detect genetic variants, the effect contributed by a single SNP-associated candidate gene may be too minor to validate. Furthermore, given the strong correlation of PHT1 to the P-acquisition phenotypes, the contribution by other single associated genes may not be easy to reveal. Therefore, the ineffectiveness of recovering the original phenotypes in the loss-of-function mutants of a single candidate gene (other than PHT1) strengthens the predominant role of PHT1 in determining the P-acquisition phenotypes.

Our findings extend current knowledge by uncovering additional genetic regulators of P acquisition and linking haplotypes to nutrient ion contents in the habitats of *A. thaliana* accessions. Furthermore, our phenotyping and analytical approaches provide a framework to systematically assess the effectiveness of the GWAS approach for studies of quantitative traits.

## MATERIALS AND METHODS

### Plant materials and growth conditions

Two panels of *A. thaliana* natural accessions (CS76309 and CS76427) used in this study were purchased from the Arabidopsis Biological Resource Center (ABRC). CS76309, which was the largest population of natural accessions available when this project was initiated, is a core set of 360 natural accessions genotyped with a 250k-SNP/ high-density tiling array by Justin Borevitz’s laboratory. (Horton et al., 2012). CS76427 is a set of 80 natural accessions sequenced using the Illumina GA platform by Detlef Weigel’s laboratory as part of the 1001 Genomes Project (Cao et al., 2011). Ninety-seven T-DNA mutants covering 43 candidate genes associated with GWAS traits were obtained from the ABRC and Nottingham Arabidopsis Stock Centre (Supplemental Table S3). Homozygous lines were isolated and analyzed under moderately low Pi conditions (50 μM KH_2_PO_4_).

Seeds were surface sterilized and germinated on agar plates with one-half strength modified Hoagland nutrient solution containing 250 μM KH_2_PO_4_, 1% sucrose, and 0.8% bactoagar (Chiou et al., 2006). Six-day-old seedlings were transferred to Pi-sufficient (+Pi) and Pi-deficient (−Pi) media supplemented with 1 mM and 50 μM KH_2_PO_4_, respectively, and grown for another 6 days. Plants were grown at 22°C under a 16-h photoperiod with cool fluorescent white light at 100–150 μE m^−2^ sec^−1^.

### Plasmid construction and plant transformation

For complementation of the *pht1;1* mutant, the open reading frame of *PHT1;1* was cloned into pCR8/GW/TOPO (Invitrogen) and then recombined into the Gateway destination vector pMDC32, in which the constitutive *35S* promoter was replaced by the *PHT1;1* promoter (3.3 kb upstream from the start codon). Modification of the sequence of *PHT1;1* was performed by site-directed mutagenesis using the Col-0 sequence as the template. The resulting constructs were designated PHT1;1^DPTSP^ and PHT1;1^NPESA^. Transgenic plants were generated using an *Agrobacterium tumefaciens* dipping procedure (Clough and Bent, 1998) with strain GV3101.

### RNA extraction and quantitative RT-PCR analysis

Total RNA from 14-day-old seedlings was isolated using TRIzol reagent (Invitrogen) followed by DNase I (Ambion) treatment. cDNA was synthesized from 1 μg total RNA using Moloney murine leukemia virus reverse transcriptase (Invitrogen) with oligo(dT) primer. Quantitative RT-PCR was performed using the Power SYBR Green PCR Master Mix Kit (Applied Biosystems) on a 7500 Fast Real-Time PCR System (Applied Biosystems) according to the manufacturer’s instructions. Relative expression levels were normalized to an internal control, *Actin8* (Supplemental Table S6). The quantitative PCR primers designed to examine transcript levels are located at the same position in the selected genes with a perfect match in each accession.

### Western blot analysis

Roots from ten 12-day-old seedlings were collected, ground in liquid nitrogen, and dissolved in protein lysis buffer containing 2% SDS, 60 mM Tris-HCl, pH 8.5, 2.5% glycerol, 0.13 mM EDTA, and 1× complete protease inhibitor (Roche). Total proteins (50 µg) were loaded onto 4% to 12% Bis-Tris sodium dodecyl sulfate-polyacrylamide gel electrophoresis (SDS-PAGE) gels (NuPAGE System) and transferred to polyvinylidene difluoride membranes. PHT1 proteins were then detected using PHT1;1/2/3 primary rabbit polyclonal antibody (Liu et al., 2011). ADP-ribosylation factor 1 (Arf-1), a Golgi marker, recognized by anti-Arf-1 antibody (Agrisera, AS08 325), was used as a loading control. The chemiluminescence signal was acquired by an UVP BioSpectrum 815 Image System (Level Biotechnology) under a linear dynamic range without saturation and quantified using ImageJ (https://imagej.nih.gov/ij/).

### Pi concentration and Pi uptake activity

Pi concentration and Pi-uptake activity were measured as described previously (Chiou et al., 2006). The shoots and roots of 12-day-old seedlings were harvested separately for the Pi concentration analysis. To measure Pi-uptake activity, five 11-day-old seedlings were first incubated in sugar-free −Pi liquid medium overnight. The next day, the Pi uptake activity was measured using whole seedlings after transferring to a medium containing 50 μM [^33^P] KH_2_PO_4_ for 2 h. The seedlings were grown under the same conditions as samples used for western blot analysis.

### GWAS

Among the 440 (360 + 80) natural accessions, we excluded those with the same ID but a different stock number (e.g., Vezzano-2 [CS76349 and CS76350]), a similar ID (e.g., Ciste-1 and Ciste-2), or seeds unavailable after propagation. We phenotyped 306 lines for Pi-uptake activity. Because some accessions could not be identified in GWA-Portal or GWAPP, we reduced the number to 180 for phenotyping PHT1;1/2/3 protein abundance.

To detect PHT1;1/2/3 protein level, we randomized the order of loading for a set of samples in one SDS-PAGE gel to avoid variation caused by transfer efficiency among replicates. In addition, an identical protein sample extracted from Col-0 accession was loaded on every SDS-PAGE gel. Each blot was also probed with anti-Arf-1 antibody to acquire the relative value of PHT1 (PHT1/Arf-1) among different accessions. This relative value was then normalized to the identical Col-0 sample between different blots to obtain the final relative PHT1;1/2/3 protein abundance in each accession from three to four biological replicates. We applied the coefficient of variation (CV) as a threshold to take the variability of biological replicates into account (Luo et al., 2019). Since CV was defined as the ratio of standard deviation (SD) to the mean, setting the filtering threshold using CV instead of SD could avoid the problem of sensitivity of SD to outliers. When CV was set to 0.3, the number of accessions was sufficient for GWAS analysis (264 and 178 *A. thaliana* accessions for Pi-uptake activity and the protein levels of PHT1;1/2/3, respectively).

The quantified data were analyzed by GWA-Portal against an imputed full sequence dataset using an accelerated mixed model (AMM), which takes kinship into account to reduce the confounding effects of population structure (Kang et al., 2008; Seren et al., 2012; Consortium, 2016). The raw values of Pi-uptake activity (Supplemental Table S1A) were subjected to analysis directly; however, square root-transformed (SQRT) values for PHT1;1/2/3 protein abundance (Supplemental Table S1B), an option implemented in GWA-Portal, were used to improve normality. For measuring the population structure of PHT1;1/2/3 protein abundance, 3,319,069 SNPs (minor allele frequency [MAF] > 0.01) were included in the datasets. For measuring Pi-uptake activity, 3,417,097 SNPs (MAF > 0.01) were included. Population structure was analyzed using TASSEL (Bradbury et al., 2007). The trait heritability was calculated from the phenotype and kinship matrix in the PyGWAS package (Provided by Ümit Seren, software engineer of GWA-portal). The LDheatmap R package was used to produce pairwise linkage disequilibrium (LD) measurements for SNPs (Shin et al., 2006).

### Dual luciferase assay

For the dual luciferase assay, the promoter of Col-0-*PHT1;1* (2 kb upstream from its start codon) was amplified and inserted into the pGreenII0800-LUC vector, which carries the firefly luciferase gene as a reporter and the renilla luciferase gene driven by the mosaic virus *35S* promoter for normalization (Hellens et al., 2005). The coding sequence of Col-0-*MYB52* fused with 10 × Myc was cloned into the pGWB520 vector. *A. thaliana* (Col-0) protoplasts were isolated and transformed with the two types of plasmids according to the method described previously (Sheen, 2001). After 12–16 h of incubation in the medium containing 50 μM KH_2_PO_4_, the protoplasts were harvested and the *PHT1;1* promoter activity was quantified with a dual-luciferase assay kit (Promega).

## Supplemental Data

**Supplemental Figure S1. Phenotypes of Col-0 seedlings grown under different concentrations of Pi.**

**(A)** Images of 12-day-old seedlings grown on media containing 250 µM Pi for 6 days, then transferred to media containing different Pi concentrations and grown for another 6 days. Bar = 1.3 cm. Fresh weights of roots **(B)** and shoots **(C)** from 10 seedlings. Pi concentration in roots **(D)** and shoots **(E)**. Error bar indicates mean ± SD (n = 3 biologically independent samples). Different letters above bars indicate statistically different groups according to one-way ANOVA followed by the least significant difference test at a probability of *P* < 0.05.

**Supplemental Figure S2. Principal component analysis (PCA) estimation of population structure in a genetic dataset.**

An imputed full sequence dataset (The 1001 Genomes Consortium, 2016) was used for PCA. PCA of PHT1;1/2/3 protein abundance **(A)** and Pi-uptake activity **(B)** (see details in “MATERIALS AND METHODS”).

**Supplemental Figure S3. A schematic map showing how a candidate gene was selected within the 10 kb region flanking a significant SNP from the association mapping of Pi-uptake activity.**

**(A)** Manhattan plot of Pi-uptake activity (also shown in Figure 1C). **(B)** Taking a significant SNP (17,400,077; −log_10_(*P*) = 7.11) on Chr 5 as an example, four genes, namely *C-NAD-MDH2* (*AT5G43330*), *PHT1;6* (*AT5G43340*), *PHT1;1* (*AT5G43350*), and *PHT1;3* (*AT5G43360*) were located within the 20-kb window centered on the SNP; each gene was scored as having one “hit”. Taking *PHT1;1* as an example, the gene body overlaps with the 20-kb windows of 61 significant SNP hits; thus, the “hit frequency” of *PHT1;1* is 61 (Supplemental Table S3). The threshold for Bonferroni correction provided by GWA-Portal is indicated on the top of the Manhattan plot **(B)** as a red dashed line.

**Supplemental Figure S4. Pi-related phenotypes of selected T-DNA mutants.**

Seedlings of 12-day-old mutant and wild-type (Col-0) plants grown under 50 μM Pi (– Pi) conditions were analyzed. **(A)** Relative protein levels of PHT1;1/2/3 in roots, n = 7 to 21 biological repeats. **(B)** Relative Pi uptake activity of whole seedlings, n= 7 to 10. The results are shown as a box and whisker diagram with dots. The bars within the box plots represent the 25th and 75th percentiles, the center line indicates the median, and the whiskers indicate the minimum and maximum. Each dot represents an individual sample. Significant differences between mutants and WT were determined by a two-sided Student’s *t*-test, and *P*-values were corrected by Benjamini Hochberg multiple testing (* Adjusted *P* < 0.05; ** Adjusted *P* < 0.01).

**Supplemental Figure S5. The transcript levels of candidate genes in Col-0 under different Pi concentrations.**

Relative mRNA levels of *PHT1;1*, *PHT1;4* **(A)** and selected candidate genes **(B)** in roots and shoots grown under sufficient Pi (+Pi; 1 mM KH_2_PO_4_, black bars) and low Pi (-Pi; 50 μM KH_2_PO_4_, white bars) conditions. Data are presented as mean ± SE (n = 4 biologically independent samples). Significance of differences was determined by two-sided Student’s *t*-test (-Pi versus +Pi treatment, ** *P* < 0.01; \**P* < 0.05).

**Supplemental Figure S6. A snapshot of the genomic region associated with the significant peak on Chr1.**

**(A)** The upper panel shows the results of a conditional GWAS of PHT1;1/2/3 protein levels for Chr1 using GWAPP. The genomic map (lower panel) shows the candidate genes surrounding the significant peak on Chr1. **(B)** All identified SNPs within the *MYB52* (*AT1G17950*) (Chr1:6,178,123, 6,178,793, and 6,178,993) gene body are in high LD (0.8 < r^2^ < 1; Supplemental Table S5B) with SNP locus 1.

**Supplemental Figure S7. LD relationships for polymorphisms within and surrounding the *PHT1;1* locus.**

**(A)** The upper panel shows the results from association mapping of PHT1;1/2/3 protein level for Chr5. The lower panel is a genomic map of the significant SNPs, including SNP locus 2, within the *PHT1;1* gene body. **(B)** SNP3 to SNP6, which are located in the second exon of the *PHT1;1* gene, result in two types of PHT1;1 protein sequence: Col-0 (NPESA) and non-Col-0 (DPTSP). **(C)** Four non-synonymous SNPs together with SNP locus 2 form four combinations of haplotypes, among which “AGAGC” (marked in blue) and “GACCT” (marked in light green) are dominant.

**Supplemental Figure S8. Expression levels of candidate genes co-localized with the significant peak identified by conditional GWAS.**

The mRNA levels of *PHT1;1*, *PHT1;2*, and *PHT1;3* **(A)** and candidate genes **(B)** in roots of different accessions. Two-way ANOVA was performed to test the epistatic relationship between two loci based on averaged mRNA levels (2^-△CT^ x 100) of *PHT1;2* **(C)** and *PHT1;3* **(D)**. The letters in blue and red shown in **(C)** and (**D)** indicate SNP locus 1 and SNP locus 2, respectively. Relative expression levels of all selected genes are shown. Data are presented as mean ± SD (n = 3 biologically independent samples).

**Supplemental Figure S9. No direct binding of MYB52 to the selected region of *PHT1;1* promoter was detected.**

**(A)** Schematic illustration of the binding sites of PHR1 (P1BS, green bars) (Rubio et al., 2001) and MYB52 (MBSI and MBSII) (Franco-Zorrilla et al., 2014) in the *PHT1;1* promoter, as analyzed by PlantPAN 2.0 (Chow et al., 2016). **(B)** Results of yeast-one-hybrid analysis conducted using the Gold Yeast One-Hybrid Library Screening System (Matchmaker). PHR1 and MYB52 were cloned into the prey vector pGADT7, and three tandem repeats of P1BS (GAATATCC) and MBSII (ATTGGC) were cloned into the pAbAi vector (Supplementary Table S4). Colony growth on leucine and uracil drop-out medium (-L-U) represents the success of co-transformation. Positive interactions were detected by selection with Aureobasidin A (AbA100 and AbA200 indicate concentrations of 100 and 200 ng/mL, respectively). Interaction between Rec-p53 and P53 served as a control.

**Supplemental Figure S10. GWAS of Pi-uptake activity among two subgroups of *A. thaliana* natural accessions based on SNP locus 2 “T” and “C” on Chr5.**

Manhattan plots from GWAS of Pi-uptake activity among 111 accessions carrying allele “T” at SNP locus 2 **(A)** or among 153 accessions carrying allele “C” at SNP locus 2 **(B)**. Thresholds for Bonferroni (red dashed line) and Benjamini Hochberg (blue dashed line) correction provided by GWA-Portal are indicated on the top of each plot. The two lines in **(B)** overlap.

**Supplemental Figure S11. Functional characterization of two forms of *PHT1;1* protein.**

**(A)** Plasma membrane localization of PHT1;1 proteins encoded by Col-0 type and non-Col-0 type *PHT1;1* transgenes. Bar = 20 μm. Relative protein levels of PHT1;1/2/3

**(B)** and Pi-uptake activities **(C)** under moderately low Pi conditions (50 μM Pi) among different plants are shown. For PHT1;1/2/3 protein level, n= 4 to 14 biologically independent samples; for uptake activity, n= 4 to 17. Numbers indicate different independent lines. Vector, pMDC32 vector in which the *35S* promoter is substituted with the *PHT1;1* promoter. The results are shown by a box and whisker diagram with dots. The bars within the box plots represent the 25th and 75th percentiles, the central line indicates the median, and the whiskers indicate the maximum and minimum. Each dot represents an individual sample. Different letters above the bars indicate statistical differences from one-way ANOVA followed by the least significant difference test at a probability of *P* < 0.05.

**Supplemental Table S1. Phenotypic values of Pi-uptake activity (A) and square root-transformed (SQRT) PHT1 protein levels (B) for GWAS analysis.**

**Supplemental Table S2. SNPs associated with Pi-uptake activity (A) and PHT1;1/2/3 protein level (B) filtered by a minor allele count (MAC) >10 and –log_10_(*P*) >4.**

**Supplemental Table S3. A list of the candidate protein-coding genes associated with Pi-uptake activity and PHT1;1/2/3 protein level located within the 10-kb regions flanking the significant SNPs.**

**Supplemental Table S4. List of the T-DNA insertional mutants examined in this work.**

**Supplemental Table S5. LD matrix of SNPs located inside *PHT1;1* (A) and near the *MYB52* (B) gene body.**

**Supplemental Table S6. List of the primer pairs used in this work.**

## ACKNOWLEDGMENTS

We acknowledge the ABRC for providing the *A. thaliana* natural accessions. We thank the Plant Tech Core Facility in the Agricultural Biotechnology Research Center of Academia Sinica for protoplast transformation and Ms. Shu-Chen Shen in the Advanced Optical Microscope Core Facility of Academia Sinica for operation of the confocal microscope. We are grateful to Ümit Seren who is the webpage manager of the GWA-Portal for helpful suggestions on GWAS analysis and technical troubleshooting. We also thank Dr. Bernhard Schäfke, a Research Assistant Professor at the Academy for Advanced Interdisciplinary Studies & Department of Biology of Southern University of Science and Technology, for constructive discussion, and Dr. Zhengrui Wang for constructive discussion and technical advice. This work was supported by the Investigator Award (AS-IA-106-L02), Academia Sinica, Taiwan.

## AUTHOR CONTRIBUTIONS

P.-S. C., C.-W. T., and T.-J. C. conceived and designed the study. P.-S. C., Y.-Y. H., and S.-F. C. performed the experiments. P.-S. C., C.-H. C., Y.-T. C., C.-W. T., and T.-J. C. analyzed the data. P.-S. C., T.-J. C., C.-H. C., Y.-T. C., and C.-W. T. wrote the manuscript.

